# Low mitochondrial diversity in native Italian pig breeds is consistent with the occurrence of strong population bottlenecks

**DOI:** 10.1101/128751

**Authors:** Joanna Kubejko, Marcel Amills, Fabio Pilla, Mariasilvia D’Andrea, Alex Clop

## Abstract

In this study, we have analysed the variation of 81 Italian pigs from the Calabrese, Casertana, Cinta Senese, Sarda and Nero Siciliano breeds as well as 54 Italian wild boars by using a dataset of mitochondrial control region sequences generated by us and others. Diversity parameters were rather low in Italian native pigs, with haplotype and nucleotide diversities ranging between 0.35-0.79 and 0.0013-0.0061, respectively. This result is consistent with the strong population bottlenecks that traditional Italian breeds have suffered due to competition with more productive foreign pig varieties. Moreover, median-joining network analysis showed that the majority of Italian pig sequences are distributed in two main clusters and that all of them belong to the E1 haplogroup. Conversely, Italian wild boars were more diverse than their domestic counterparts and they harboured the E1 and E2 haplogroups. The absence of the E2 haplogroup in Italian pigs and its moderate frequency in wild boars might suggest that this haplogroup was rare at the time that wild boars were domesticated in Italy.

---

At the beginning of the 20^th^ century, there were 21 traditional pig breeds in Italy (Mascheroni 1927), but the majority of them disappeared as a consequence of changes in the use of land, the industrialisation of pig production and the massive use of highly improved imported breeds that occurred from the nineteen fifties onwards. For instance, The Friuli Black breed was crossed with Edelschwein and Large White pigs to an extent that it became practically extinct in that decade (Porter et al. 2016). In Northern Italy, many ancient breeds have been definitively lost (*e.g.*Valtellina, Garlasco, Borghigiana and Modenese), while in the South the number of breeds that have survived is greater though several of them have an endangered or critical status (Porter et al. 2016). In recent times, the interest about local breeds has grown because they can yield highly valued products (*e.g*. cured ham and sausage) and they are a reservoir of genetic diversity (D’Andrea *et al*., 2008; D’Andrea *et al*., 2011).

Population sizes of the few Italian pigs breeds (Calabrese, Casertana, Cinta Senese, Mora Romagnola, Nero Siciliano and Sarda) that have managed to survive are still rather small (Franci and Pugliese 2007) and their mitochondrial diversity appears to be reduced (Scandura et al. 2008). From a phenotypic point of view, these breeds have black or grey coats, longish snouts and ears covering their eyes (Porter et al. 2016). In this work, we aimed to investigate the consequences of population decline on the mitochondrial variation of several indigenous Italian pig breeds. Moreover, we were interested in comparing the diversity of Italian swine and wild boars because the latter are known to harbour mitochondrial haplotypes that have been found only in Italy (Giuffra et al. 2000, Larson et al. 2005).

Genomic DNA was extracted from hair, skin or skeletal muscles from 35 Nero Siciliano, Calabrese and Casertana pigs using standard protocols. The mitochondrial control region was amplified by following the protocols reported by Noce et al. (2015). Amplified products were purified with the ExoSAP kit (Thermo Scientific Company) and sequenced on both directions with the BigDye Terminator v3.1 Cycle Sequencing kit (Applied Biosystems) and primers reported by Noce et al. (2015). Sequencing reactions were electrophoresed in an ABI 3730 DNA analyzer (Life Technologies). Control region sequences generated in this way were deposited in GenBank (accession numbers from KY942150 to KY942183). In our analysis, we have also used a data set of 148 sequences retrieved from GenBank and corresponding to Italian pigs and wild boars as well as to Chinese and European wild boars (**Supplementary Table 1**).

A median-joining network was built with the Network 4.6. software (Bandelt et al. 1999) following the procedures reported by Noce et al. (2015). This analysis indicated that most of Italian domestic pigs, with the exception of a few Nero Siciliano individuals, group in two main clusters (**Figure 1**). We also estimated several diversity parameters with the DnaSP 5.10 software (Librado and Rozas 2009). Haplotype and nucleotide diversities of Italian pig breeds ranged between 0.35-0.79 and 0.0013-0.0061, respectively (**Table 1**), a result that is consistent with. Scandura et al. (2008). Moreover, the nucleotide diversity of Italian wild boars was higher than that observed in native pigs (**Table 1**). Tajima D-statistics calculated in the four Italian pig populations were positive but not significant (data not shown), indicating the absence of a relevant departure from the neutrality expectation. Guastella et al. (2010) analysed the variation of Nero Siciliano pigs with a panel of microsatellite markers and found that the expected heterozygosity was 0.70, which is a relatively high value. However, microsatellites included in diversity panels are usually chosen because of their high variability, which typically inflates diversity estimates. Besides, reductions in population size are expected to have more drastic effects on the mitochondrial than on the autosomal variation because of the lower effective size of the former. We did not detect Asian haplotypes in Italian swine, a result that is consistent with data presented by Clop et al. (2003) indicating that many European local breeds were spared from the process of introgression with Chinese sows that took place in the United Kingdom during the 18-19^th^ centuries.

**Table 1.** Nucleotide and haplotype diversity of the mitochondrial control region in Italian wild boars and pigs (Sarda pigs have not been included due to their low sample size).

| Population | N | Nucleotide diversity | Haplotype diversity |
| --- | --- | --- | --- |
| Italian wild boar | 54 | $0.0112 \pm 0.0010$ | $0.748 \pm 0.047$ |
| Calabrese | 19 | $0.0037 \pm 0.0004$ | $0.743 \pm 0.062$ |
| Casertana | 26 | $0.0028 \pm 0.0007$ | $0.369 \pm 0.091$ |
| Cinta Senese | 10 | $0.0013 \pm 0.0006$ | $0.356 \pm 0.159$ |
| Nero Siciliano | 21 | $0.0061 \pm 0.0005$ | $0.795 \pm 0.051$ |

**Figure 1.**
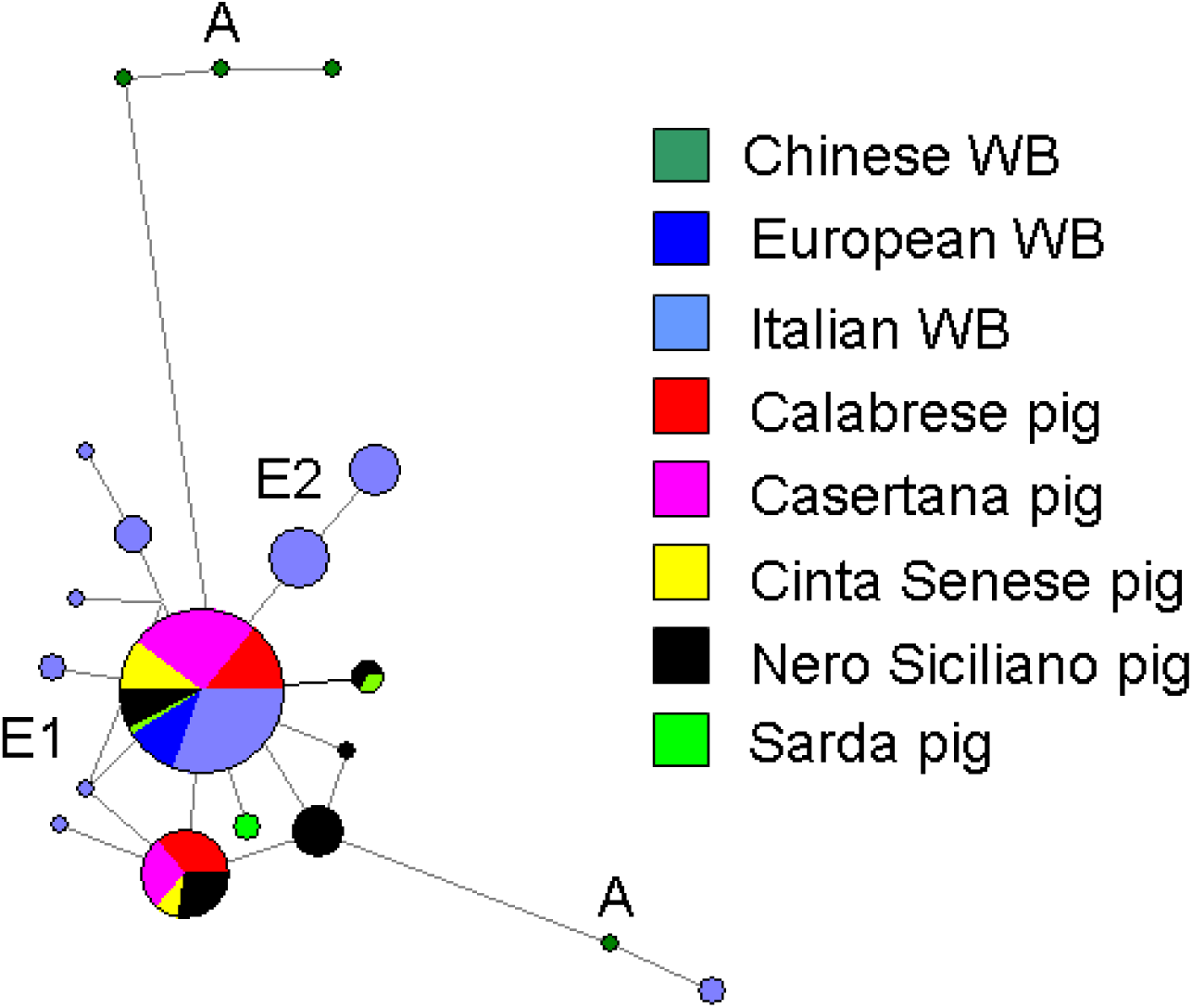
Median-joining network of mitochondrial control region sequences from Italian wild boars and domestic pigs. Asian and European 1 and 2 haplogroups are marked as A, E1 and E2, respectively. A few sequences from Chinese, Austrian and Belgian wild boars have been also included as a reference.

The low variation of Italian pigs is compatible with a sustained demographic recession and the occurrence of genetic bottlenecks. According to Franci and Pugliese (2007), the population sizes of Cinta Senese, Casertana, Calabrese and Nero Siciliano pigs recorded in the Pedigree Register of 2006 were 1395, 84, 84 and 169, respectively. However, for several of these breeds (*e.g.*Nero Siciliano) the numbers might be higher due to the difficulty of registering individuals bred in free-range systems (Franci and Pugliese 2007). Importantly, demographic declines have been often sharp and sudden. For instance, by the mid-fifties there were 160,000 Cinta Senese pigs, but a decade later, this breed was at the verge of extinction due to the massive importation of highly improved varieties and in 1986 only 81 purebred sows and 3 boars were registered (Scali et al. 2012). With the aim of preserving this breed, during the seventies and eighties it was extensively crossed with Landrace and Large White pigs, thus provoking the dilution of the original genetic background. However, we have not found any evidence of the segregation of Asian mitochondrial variants in Cinta Senese pigs despite the fact that these variants are fairly common in Large White swine (Fang and Andersson 2006).

Giuffra et al. (2000) was the first to report the existence of a European mitochondrial haplogroup E2 that thus far has been only found in Italian wild boars (Larson et al. 2005). Larson et al. (2005) renamed this haplogroup as D4, and hypothesized that it may have an early Italian origin. The median-joining network shown in **Figure 1** indicates that the E2/D4 haplogroup does not segregate in any of the Italian pigs. This result corroborates data published by Scandura et al. (2008), who analysed the mitochondrial variation of 98 wild boars and 47 swine from Italy and demonstrated that all variants detected in domestic pigs belong to the E1 haplogroup, whilst the E2 segregates exclusively in wild boars at an approximate frequency of 0.36. Given its frequency in wild boars, the complete absence of the E2 haplogroup in Italian native pig breeds is paradoxical. One possible interpretation would be that the E2 haplogroup had a low frequency in ancient times, when wild boars began to be locally domesticated in Italy, but it expanded recently in the Italian wild boar population. Between the 17^th^ and 19^th^ centuries, wild boars disappeared from many parts of Italy such as Trentino, Liguria, Friuli and Romagnola (Apollonio et al. 1988), and the beginning of the 20^th^ century wild boar distribution was restricted to Sardinia and several central and southern areas of Italy (Ghigi 1911). After the Second World War, the Italian wild boar populations began to grow exponentially (Scandura et al. 2008), providing an opportunity for the stochastic and sharp increase in the frequency of variants that were not abundant before the bottleneck (Zhang et al. 2004). Alternatively, Italian wild boars carrying the E2 haplogroup did not contribute to the initial pool of domestic pigs as a consequence of restricted gene flow (Veličković et al 2016).

The main conclusion of the current work is that Italian local pig breeds display low levels of mitochondrial diversity, probably as a consequence of strong bottlenecks that took place in recent and ancient times. According to FAO, one third of animal breeds face extinction, causing the loss of genetic variants that cannot be replaced (http://www.fao.org/News/2000/001201-e.htm). In Italy, there were 21 porcine breeds at the beginning of the 20^th^ century but only six have managed to survive up to present time (Franci and Pugliese 2007). Our results support the notion that the genetic conservation of these six breeds should be focused on implementing adequate mating plans aiming to maximize variability thus avoiding the detrimental effects of inbreeding (Franci and Pugliese 2007).

## Acknowledgments

This research was partially funded with projects AGL2013-48742-C2-1-R and AGL2013-44978-R granted by the Spanish Ministry of Economy and Competitiveness. Alex Clop acknowledges the Ramon y Cajal fellowship awarded by the Spanish Ministry of Economy and Competitiveness (RYC-2011-07763). We also acknowledge the support of the Spanish Ministry of Economy and Competitiveness for the *Center of Excellence Severo Ochoa 2016-2019* (SEV-2015-0533) grant awarded to the Center for Research in Agricultural Genomics. Thanks also to the CERCA Programme of the Generalitat de Catalunya.

---

**Supplementary Table 1.** List of pig and wild boar mitochondrial control region sequences retrieved from GenBank (http://www.ncbi.nlm.nih.gov/genbank).

| **Population** | **GenBank acc. number** |
| --- | --- |
| Wild boar from Forli | EU362417- EU362421 |
| Wild boar from Siena | EU362427- EU362431 |
| Wild boar from Salerno | EU362435 – EU362439 |
| Wild boar from Florence | EU362445 – EU362449<br>EU362453 – EU362454<br>AY884724 |
| Wild boar from San Rossore | EU362466 – EU362469<br>EU362475 |
| Wild boar from Castel Porziano | EU362476 - EU362479 |
| Wild boar from Arezzo | AM748931<br>AM748934<br>AM744976<br>AM748936 |
| Wild boar from Bergamo | AM748930<br>AM748937<br>AM748933<br>AM748935 |
| Wild boar from Maremma | EU362455 - EU362457<br>EU362465<br>AY884716 - AY884722 |
| Wild boar from Parma | AM773230 |
| Wild boar from Italy | AB015094 |
| Calabrese pig | EU362569 - EU362573 |
| Casertana pig | EU362575 - EU362578<br>EU362580 - EU362585 |
| Cinta Senese pig | EU362559 - EU362568 |
| Nero Siciliano pig | EU362586 - EU362600 |
| Sarda pig | EU362554 - EU362558 |
| Wild boar from China | AY884610<br>AY884639<br>AY884640<br>AY884683 |
| Wild boar from Belgium | DQ379234 - DQ379237<br>DQ379240 |
| Wild boar from Austria | HM747196 |
|  | HM747197<br>NM214041 |

